# Segmentor: A tool for manual refinement of 3D microscopy annotations

**DOI:** 10.1101/2021.01.25.428119

**Authors:** David Borland, Carolyn M. McCormick, Niyanta K. Patel, Oleh Krupa, Jessica T. Mory, Alvaro A. Beltran, Tala M. Farah, Carla F. Escobar-Tomlienovich, Sydney S. Olson, Minjeong Kim, Guorong Wu, Jason L. Stein

**Affiliations:** RENCI, University of North Carolina at Chapel Hill, Chapel Hill, NC 27599, USA; UNC Neuroscience Center, University of North Carolina at Chapel Hill, Chapel Hill, NC 27599, USA; Department of Genetics, University of North Carolina at Chapel Hill, Chapel Hill, NC 27599, USA; Department of Computer Science, University of North Carolina at Greensboro, NC 27412, USA; Department of Psychiatry, University of North Carolina at Chapel Hill, Chapel Hill, NC 27599, USA; Department of Computer Science, University of North Carolina at Chapel Hill, Chapel Hill, NC 27599, USA

**Author notes:** Correspondence to Guorong Wu: 334 Emergency Room Drive, 343 Medical Wing C, Chapel Hill, NC 27599, USA, Correspondence to Jason Stein: 116 Manning Drive, CB# 7250, University of North Carolina at Chapel Hill, Chapel Hill, NC 27599, USA. These authors jointly supervised the work (Guorong Wu, Jason L. Stein).

**Keywords:** tissue clearing, light sheet microscopy, deep learning, image segmentation, manual annotation

## Abstract

**Background:** Recent advances in tissue clearing techniques, combined with high-speed image acquisition through light sheet microscopy, enable rapid three-dimensional (3D) imaging of biological specimens, such as whole mouse brains, in a matter of hours. Quantitative analysis of such 3D images can help us understand how changes in brain structure lead to differences in behavior or cognition, but distinguishing features of interest, such as nuclei, from background can be challenging. Recent deep learning-based nuclear segmentation algorithms show great promise for automated segmentation, but require large numbers of manually and accurately labeled nuclei as training data.

**Results:** We present Segmentor, an open-source tool for reliable, efficient, and user-friendly manual annotation and refinement of objects (e.g., nuclei) within 3D light sheet microscopy images. Segmentor employs a hybrid 2D-3D approach for visualizing and segmenting objects and contains features for automatic region splitting, designed specifically for streamlining the process of 3D segmentation of nuclei. We show that editing simultaneously in 2D and 3D using Segmentor significantly decreases time spent on manual annotations without affecting accuracy.

**Conclusions:** Segmentor is a tool for increased efficiency of manual annotation and refinement of 3D objects that can be used to train deep learning segmentation algorithms, and is available at https://www.nucleininja.org/ and https://github.com/RENCI/Segmentor.

## Background

The structure of the brain provides the machinery that enables behavior and cognition. The human brain is extremely complex, comprising ∼170 billion cells, of which ∼86 billion are neurons [1]. The mouse brain, a common model system used to study brain-behavior relationships, is much smaller yet still has ∼109 million cells, ∼70 million of which are neurons [1]. By mapping the location of these many brain cells, classifying them into types based on the expression of marker genes, and determining how cell type proportions and locations are altered by mutations or environmental factors, we can understand how changes in brain structure lead to changes in behavior and/or cognition.

In order to map cell types within intact brains, a number of tissue clearing techniques for making the brain transparent were recently introduced [2, 3]. Combined with high-speed image acquisition through light sheet microscopy, the full 3D extent of adult mouse brain specimens can be imaged at micron resolution in a matter of hours [4–7].

Currently, these large-scale microscopy images are often used for qualitative visualization rather than quantitative evaluation of brain structure, thus potentially overlooking key spatial information that may influence structure-function relationships for behavior and cognition. In order to quantify objects within annotated regions of the images, we need to distinguish morphological objects of interest (e.g., nuclei) from background [8]. Existing programs that perform object segmentation in cleared samples from tissue (for example, ClearMap [9], CUBIC [10]) or organoids [11] work well for cases with unambiguous morphological characteristics. However, for cases in which morphological objects are densely packed, nuclei segmentation results are less accurate using current computational tools, which obfuscates brain structure quantifications and comparisons. Recent deep learning-based nuclear segmentation algorithms such as multi-level convolutional neural networks show great promise for more accurately identifying each individual nucleus [12– 15]. When colocalized with immunolabeling, nuclear segmentation additionally enables counting individual cell types. Present learning-based methods require two sets of manually labeled ‘gold standards’: (1) a large number of training objects to learn the morphometrical appearance of nuclei in the context of various backgrounds, and (2) independent benchmark datasets for evaluating the accuracy of automated segmentation results.

Gold standard datasets are derived from manual labels by trained and reliable raters. Manual labeling is both time-consuming and difficult because of ambiguities in nuclear boundaries and the difficulty of labeling 3D structures on a 2D screen. A few tools have been developed for manual labeling of objects in 2D [16] and 3D [17–20] images, including labeling in virtual reality environments [21]. Existing annotation software often implement automatic contour completion using deformable model techniques such as active contour [22]. This feature is useful when nuclei are spatially separate from each other, but is less powerful for annotating densely-packed nuclei, where the boundary between multiple nuclei has lower contrast. Additionally, existing tools are not optimized for editing large images with high visual complexity where densely packed nuclei within a 3D scene can obscure the region being edited, and, at present, do not provide methods for semi-automated correction of common automated segmentation errors (e.g., incorrectly merged or split nuclear boundaries) that lower throughput of manual refinement.

Here, we present Segmentor, an open-source tool for reliable, efficient, and user-friendly manual annotation and refinement of objects (e.g., nuclei) within 3D light sheet microscopy images. This tool enables automated pre-segmentation of nuclei, refinement of objects in 2D and 3D, visualization of each individual nucleus in a dense field, and semi-automated splitting and merging operations, among many other features. This tool has been used by 10 individuals to achieve reliable segmentation and labeling of thousands of nuclei. We show that editing simultaneously in both 2D and 3D significantly decreases labeling time, without impacting accuracy. Software releases of this tool and example images are available at https://www.nucleininja.org/, and source code and documentation are available at https://github.com/RENCI/Segmentor. We expect that increasing the number of manually labeled nuclei in 3D microscopy images through this user- efficient tool will help implement fully automated nuclear recognition by incorporating deep learning.

## Implementation

### Software

The Segmentor tool was developed in C++ using open-source cross-platform libraries, including VTK [23] and Qt [24]. 3D image volumes and segmentation data can be loaded in TIFF, NIfTI, or VTI format. In order to increase efficiency, the tool is primarily designed for manually refining existing annotations rather than beginning annotations completely anew. The user can load initial segmentation data generated by a tool external to Segmentor (e.g., NuMorph [12], CUBIC [10], or ClearMap [9]), or generate an initial global intensity threshold-based segmentation [25] from within Segmentor. The interface consists of panels with 2D (right panel) and 3D (left panel) views and a region table (**Figure 1**). The 2D view consists of a single slice through the volume, and enables the user to see both the voxel intensities, and 2D visualizations of the segmented regions. The 3D view enables the user to see 3D surfaces of the segmented regions, and inspect them for non-uniform morphology that is difficult to visualize using only the 2D view. The 3D and 2D views are synchronized, such that navigating (i.e., rotating, translating, or zooming) in one view also updates the other view. The hybrid 2D/3D visualization and editing capabilities are important as each view is useful for different aspects of the annotation procedure, e.g., the 2D view is useful for manually selecting voxels based on image intensity, whereas the 3D view is useful for identifying incorrectly segmented regions of densely packed nuclei.

**Figure 1:**
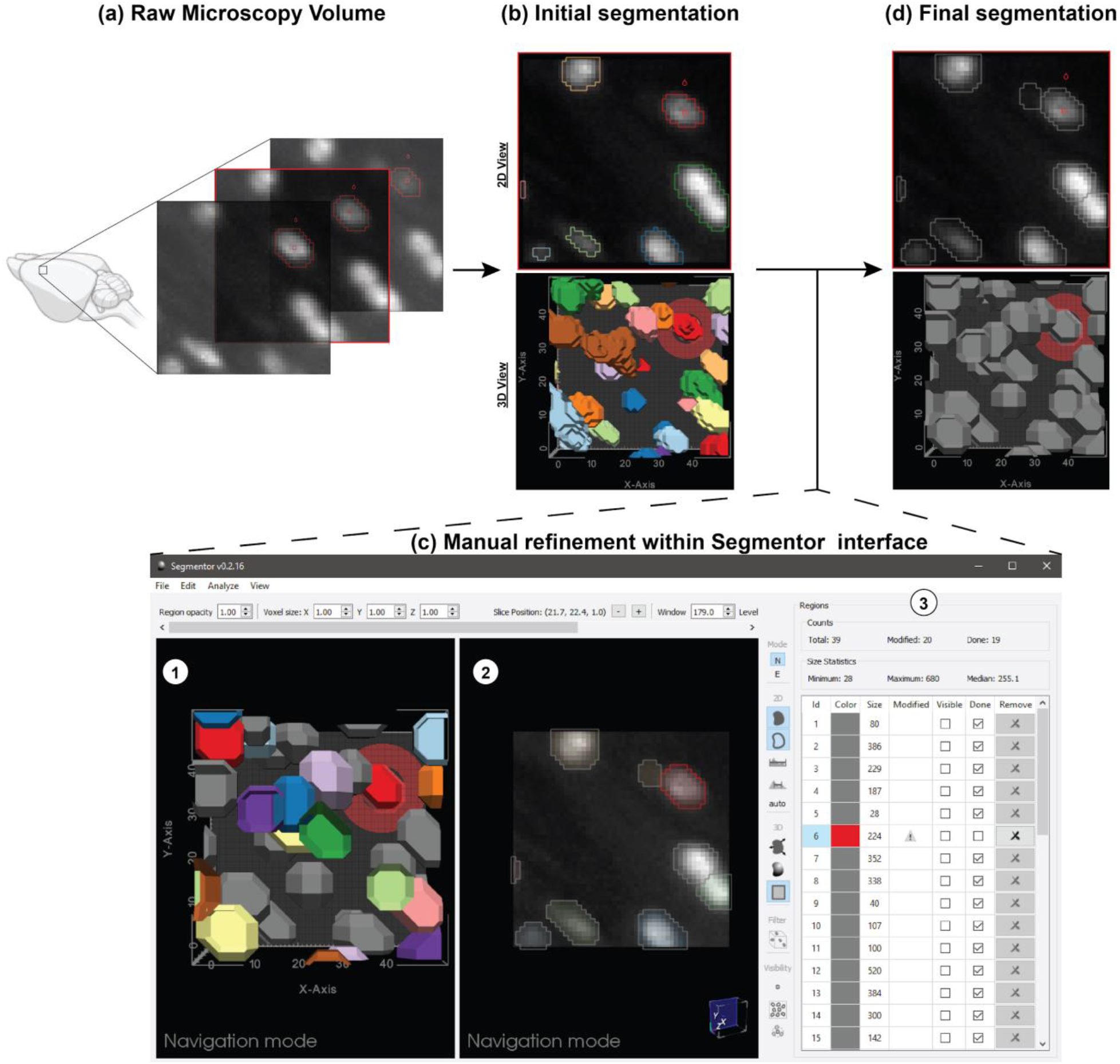
Demonstration of Segmentor software for nuclear refinement. (a) Raw microscopy volumes of the brain are loaded into the software. (b) Segmentor provides an initial segmentation of nuclei within the image (alternatively, pre-segmentations from other programs can be loaded). (c) The segmented images are manually refined within Segmentor using (1) the 3D visualization of segmented nuclei and (2) the 2D slices. (3) The region table enables the user to track progress during segmentation. (d) Finally, the manually refined image that can be used as gold standard input to deep learning programs is shown (grey regions indicate those the user has marked as completed). Image made in part using BioRender.

### Visualization features

The 2D view provides outline and filled representations of the segmented regions, and window/level controls for the voxel intensities. The 3D view has controls for smooth shading and surface smoothing. The user can also toggle a representation of the current 2D slice plane in the 3D view. Edits made in either view are immediately updated in the other view. To reduce visual clutter in the 3D view, various modes are available to filter the currently visible regions to: 1) current slice plane, 2) the currently selected region and close neighbors, and 3) the currently selected region only.

### Editing features

Various editing features are provided. Most operations can be applied in either the 2D or 3D view, although certain features are only applied in the 2D X-Y plane due to the improved resolution in that plane for most microscopy volumes. Standard editing features include voxel-level painting and erasing of the currently selected region. The user can select a brush radius, applied in the X-Y plane, for these edits.

In addition to these standard editing features, more advanced features are also provided. The user can apply a constrained region growing or shrinking operation in the X-Y plane by selecting a voxel outside (growing) or inside (shrinking) the current region. For region growing, all voxels with an intensity equal to or higher than the selected voxel that are reachable from the current region, and no farther than the selected voxel, are added to the region. This is similar to a dilation of the current region, but only including voxels with intensities greater than or equal to the current region. Region shrinking works similarly, but removes voxels with intensities less than or equal to the selected voxel.

Common segmentation problems from automatic methods include divided nuclei, where more than one region is present within a single nucleus, and joined regions, where multiple nuclei are incorrectly included as the same region. Semi-automated methods are provided for correcting these issues. To fix divided nuclei, the user can select any region to merge with the current region by reassigning the voxel labels. Splitting joined regions is more challenging (**Figure 2**). We employ an intensity threshold method: using the 2D and 3D view the user determines how many nuclei are in the current region that should be separated. After specifying this number, a fully-automated approach is applied. An increasing intensity threshold is repeatedly applied to the voxels in the region. As the intensity increases, the region is typically broken up into smaller regions. The threshold resulting in the specified number of regions (via connected component analysis) with the largest volume for the smallest of the three regions (making the method less sensitive to noise) is used to define seed regions (intensities are typically higher toward the center of the nuclei). Each seed region is then successively grown similarly to the region growing method described above, by stepping the region growing intensity down from the seed region threshold, constraining the growing to a 1-voxel radius at each step, and to the original region voxels. After splitting, the user can perform any necessary adjustments using the other editing features.

**Figure 2:**
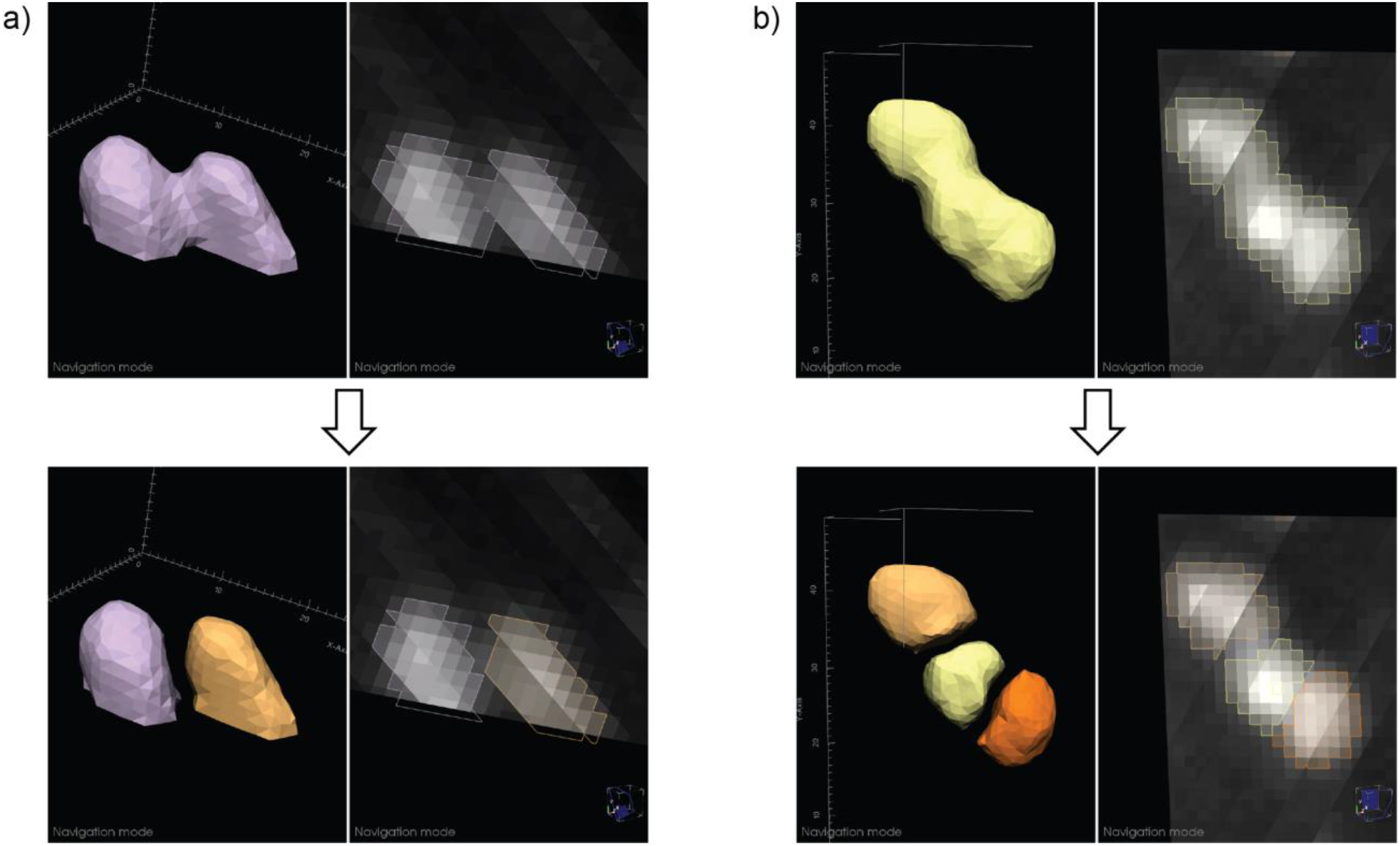
Examples of automated nuclear splitting within Segmentor. (a) An incorrectly joined region is shown (top), which after visual inspection is determined to represent two nuclei. After the user specifies that there are two nuclei in the joined region, the automated splitting function result is shown (bottom). (b) Similar to (a), but three nuclei are incorrectly joined (top) and the automated result is shown (bottom).

### Region table

To help the user manage the complexity of segmenting many nuclei in a given volume (e.g. ∼460 nuclei are found within an image volume of 152.5 μm × 152.5 μm × 248 μm of the cortex), a region table provides information on each segmentation region, including label color, size (in voxels), modified status (whether the label has been modified since the last save), and done status (whether the user considers segmentation complete for that region). The user can sort by label, size, or status, and select any region to zoom in on that region in the 2D and 3D views. The user can mark any region as done to keep track of their progress. Such regions will be greyed out in the other views. Modified and done statuses are stored in a separate JSON metadata file stored with the segmentation data.

### Typical workflow

All users undergo an initial training period in which they receive the same standardized training image containing 39 nuclei. Each user then generates an initial automated segmentation, which s/he manually edits. Labeling reliability is then iteratively assessed by comparing segmentations to those of an experienced rater (CMM) until a Dice score [26] of ≥ 0.85 is achieved and label counts are within ± 1 nucleus of the ‘gold standard’ training segmentation (i.e., 39 ± 1 nuclei).

### Case study

To quantify efficiency and accuracy of manual labeling in 2D+3D as compared to 2D alone, two raters (CMM, NKP) manually refined a series of four images using either ‘2D only’ or ‘2D+3D’ visualizations. Both raters used Segmentor v0.2.11 (Windows version) and achieved reliability on a separate standardized image prior to beginning the case study. One rater (NKP) was assigned these 4 images balanced with respect to the order of ‘2D only’ or ‘2D+3D’, to minimize ordering bias. This rater alternated between ‘2D only’ and ‘2D+3D’ using a toggle- enabled feature in Segmentor’s interface designed to hide the 3D visualization. In total, this rater completed 2 manual refinements on each of 4 images (i.e., labeling the same image twice per visualization modality). The other rater (CMM) edited each of the four images in ‘2D+3D’ only for accuracy assessment. Both tracers recorded the time to completion using the freely available Clockify application. Manually refined annotations were compared between raters for accuracy (Dice score, DSC) and differences in time and accuracy between ‘2D only’ and ‘2D+3D’ were evaluated using a paired t-test.

### Image acquisition

Images used as input to Segmentor were acquired from iDISCO+ [9] tissue clearing of P15 C57Bl/6J mice. Nuclei were labeled with TO-PRO-3 and imaged on a lightsheet microscope (Ultramicroscope II from LaVision Biotec) at a final resolution of 0.75 μm × 0.75 μm × 2.50 μm. Blocks from the cortex were used for labeling with Segmentor. Further details about image acquisition can be found in [12].

### User survey

Segmentor usability feedback was collected from six participants in a 28-question survey (24 Likert scale questions on a 7-point scale, in which ‘1’ means ‘not useful’ and ‘7’ means ‘extremely useful,’ followed by 4 open-ended questions). Qualtrics^XM^ was used to distribute the survey and analyze participant results [see Survey Results].

## Results

Ten users have used Segmentor for manual refinement of 3D microscopy volumes. Segmentations from one expert user were defined as the gold standard and results from every other user were compared to this segmentation via Dice score and nuclei counting to assess reliability. After 5 iterations, a Dice score of ≥ 0.85 was achieved by each user (average final Dice score=0.876).

To test whether simultaneous visualizations of 2D and 3D segmentations led to increased efficiency or accuracy, we designed a case study in which one user labeled nuclei using either the 2D view alone or both the 2D and 3D view in 4 images containing on average 39 nuclei. A separate expert user annotated the same images with both the 2D and 3D view to serve as gold standard for accuracy comparisons. The use of both 2D and 3D led to a 1.8-fold reduction in the amount of time needed for segmentation (2D: 554 +/- 15 min; 2D+3D: 304 +/- 21 min; p=0.00027; **Figure 3**). Using both the 2D and 3D view, manual annotation of the full 3D extent of a nucleus takes approximately 8 minutes. However, we found that use of both 2D and 3D views was not associated with differences in annotation accuracy relative to the gold standard rater (mean Dice score for 2D: 0.82 +/- 0.024; mean Dice score for 2D+3D: 0.81 +/- 0.023; p=0.86; **Figure 3**). These findings indicate that combined use of the 2D and 3D views increase speed for manual refinements without sacrificing accuracy in segmentation.

**Figure 3:**
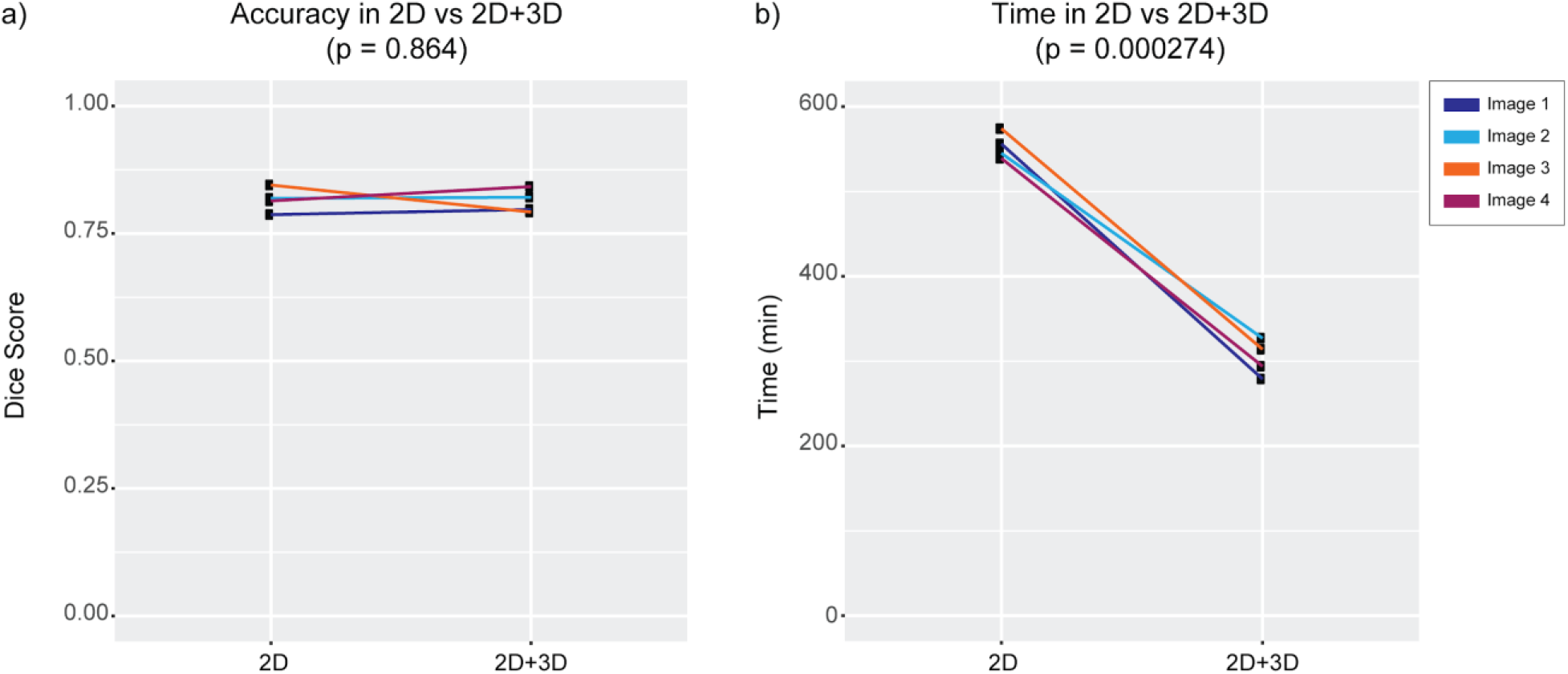
Case study to determine accuracy and efficiency of manual refinement when visualizing 2D and 3D nuclei. (a) Dice score measuring accuracy relative to an expert rater for either the labels only from the 2D segmentations or from 2D + 3D segmentations. (b) Time comparison between 2D vs 2D+3D.

The user survey corroborated with case study results, as 2D and 3D views were both found to be useful. Questions focused on the usefulness of editing segmentations in 2D and 3D received respective means of 6.33 (Q1) and 6.83 (Q2) on a 7-point Likert scale, and questions focused on the usefulness of 2D and 3D visualizations received respective means of 5.5 (Q3) and 7.0 (Q4). The region splitting feature was also confirmed to be useful, with a mean of 6.67 (Q6), and questions addressing features related to the region table had an overall mean of 6.63 (Q11-14). Visualizing non-axis-aligned slices in the 2D view supports synchronization of the 2D and 3D views, but scores on the utility of this feature varied, with a mean value of 3.33 and a standard deviation of 2.43 (Q10), perhaps due to artifacts caused by voxel anisotropy. Future work will explore more flexible coupling of the 2D and 3D views to more effectively utilize the strengths of each view.

## Discussion

A user-friendly tool for manual delineation of nuclei in 3D image volumes will greatly accelerate training of automated recognition algorithms necessary to quantify nuclei in tissue cleared images of the brain. Here, we present Segmentor as a tool to make manual 3D segmentations easier and more efficient. Segmentor has been tested and iteratively updated based on the feedback of 10 users. Segmentor provides new features that allow the user to parse relevant information and navigate in dense images, automatically split or merge nuclei, keep track of progress during segmentation, and efficiently use both 2D and 3D visual information. While we have demonstrated use-cases for nuclear segmentation, Segmentor also can be used to annotate any other features found in 3D microscopy images.

Here, we focus on identifying the borders of the 3D extent of the nucleus rather than using a marker to label one voxel within the nucleus. Though counting applications only require one voxel (or crosshair) within a nucleus to be labeled, labeling the boundaries of nuclei enables measurements of nuclear shape, facilitates more accurate colocalizations with markers across channels, and allows for evaluation of precision and recall by determining whether an automated segmentation lies within the boundaries of the manually defined nucleus. We also believe that the added information of the nuclear boundaries will provide more useful heuristics to deep learning approaches about contextual features that distinguish the nucleus from background and possibly other (touching) nuclei [14].

How many manually annotated nuclei are sufficient for training a successful image segmentation tool using deep learning methods? In recent work [14], 80,692 manually labeled nuclei (from 1,102 images) were used to train a highly accurate 2D segmentation method [27]. Learning 3D nuclei segmentation is more challenging than its 2D counterpart, so it is necessary to develop more complex neural networks (with more parameters), which require larger numbers of training samples for fine tuning the network parameters. Each 3D nucleus is composed of ∼5 slices of 2D segmentations at the image resolution used here. Thus, our goal is to acquire ∼20,000 high-quality manual 3D nuclei annotations using our Segmentor software (comprising ∼100,000 2D masks), which will be used to train, validate, and test our neural network in a 10-fold cross validation manner.

We show a case study that visualization in both 2D and 3D views increases efficiency without impacting accuracy, while significantly reducing tracing time. Because a large number of training samples are needed to train a deep learning-based segmentation model, suggested improvement of manual labeling efficiency will greatly contribute to the performance of automated segmentation software.

Finally, the current approach involves the segmentation of a full 3D image containing 40-400 cells, which still can take 5 to 50 hours of manual effort per user. We expect that as automated pre-segmentations are improved through additional manual training, time for manual refinement will decrease because less manual refinements will be required. Additionally, we expect that by chunking these segmentation tasks into smaller units of single cells or clumps of cells, more users can participate in segmentation simultaneously with less overall time commitment. This would allow annotations at a massive scale, through a larger scale citizen science approach.

## Conclusions

Segmentor is a freely available software package that increases efficiency of manual refinement in 3D microscopy images. We expect that use of this software will greatly increase the number of training samples and thereby result in higher accuracy of learning-based automated segmentation algorithms, enabling the efficient quantification of brain structural differences at cellular resolution.

## Supporting information

Survey Results

## Availability and requirements

Project name: Segmentor

Project home page: https://www.nucleininja.org/

Operating systems: Windows, Mac Programming language: C++

License: MIT

Any restrictions to use by non-academics: No restrictions

### List of abbreviations

2D: two-dimensional
3D: three-dimensional
DSC: Dice score
μm: micrometers
P15: postnatal day 15
VTK: Visualization Toolkit

## Declarations

### Ethics approval and consent to participate

The user survey was determined exempt by the UNC Office of Human Research Ethics (IRB 20- 3341).

### Consent for publication

Not applicable.

### Availability of data and materials

The training image dataset analysed during the current study is available from the project webpage: https://www.nucleininja.org/download

## Authors’ contributions

DB developed the Segmentor tool. GW and JLS initiated and supervised the project and provided funding. MK maintains the Nuclei Ninja website. JLS and CMM designed the case study. OK acquire all images used for the case study. CMM and NKP performed all segmentations for the case study. CMM performed all analyses for the case study. DB and CMM designed the user survey. CMM analyzed the survey results. JTM, AAB, TMF, CFE, and SSO performed manual segmentations and tested the software. All authors contributed to drafting and reviewing the manuscript for final approval.

## Acknowledgments

We thank Pablo Ariel of the Microscopy Services Laboratory and Michelle Itano of the Neuroscience Microscopy Core for assisting in sample imaging. We thank Matthew C. Bailey, Felix A. Kyere, Karen Huang, Jade A. Hardwick, Suh Hyun Lee, Jordan M. Valone, and Shivam Bharghava for contributing manual annotations and testing the software.

## References

1. Herculano-Houzel S, Mota B, Lent R. Cellular scaling rules for rodent brains. Proc Natl Acad Sci U S A. 2006;103:12138–43.

2. Richardson DS, Lichtman JW. Clarifying Tissue Clearing. Cell. 2015;162:246–57.

3. Ueda HR, Dodt H-U, Osten P, Economo MN, Chandrashekar J, Keller PJ. Whole-Brain Profiling of Cells and Circuits in Mammals by Tissue Clearing and Light-Sheet Microscopy. Neuron. 2020;106:369–87.

4. Becker K, Jährling N, Kramer ER, Schnorrer F, Dodt H-U. Ultramicroscopy: 3D reconstruction of large microscopical specimens. J Biophotonics. 2008;1:36–42.

5. Reynaud EG, Peychl J, Huisken J, Tomancak P. Guide to light-sheet microscopy for adventurous biologists. Nat Methods. 2015;12:30–4.

6. Tomer R, Khairy K, Keller PJ. Light sheet microscopy in cell biology. Methods Mol Biol. 2013;931:123–37.

7. Winter PW, Shroff H. Faster fluorescence microscopy: advances in high speed biological imaging. Curr Opin Chem Biol. 2014;20:46–53.

8. Caicedo JC, Roth J, Goodman A, Becker T, Karhohs KW, McQuin C, et al. Evaluation of Deep Learning Strategies for Nucleus Segmentation in Fluorescence Images. bioRxiv. 2018;:335216. doi :10.1101/335216.

9. Renier N, Adams EL, Kirst C, Wu Z, Azevedo R, Kohl J, et al. Mapping of Brain Activity by Automated Volume Analysis of Immediate Early Genes. Cell. 2016;165:1789–802.

10. Murakami TC, Mano T, Saikawa S, Horiguchi SA, Shigeta D, Baba K, et al. A three-dimensional single-cell-resolution whole-brain atlas using CUBIC-X expansion microscopy and tissue clearing. Nat Neurosci. 2018;21:625–37.

11. Piccinini F, Balassa T, Carbonaro A, Diosdi A, Toth T, Moshkov N, et al. Software tools for 3D nuclei segmentation and quantitative analysis in multicellular aggregates. Comput Struct Biotechnol J. 2020;18:1287–300.

12. Krupa O, Fragola G, Hadden-Ford E, Mory JT, Liu T, Humphrey Z, et al. NuMorph: tools for cellular phenotyping in tissue cleared whole brain images. Cold Spring Harbor Laboratory. 2020;:2020.09.11.293399. doi :10.1101/2020.09.11.293399.

13. Tokuoka Y, Yamada TG, Mashiko D, Ikeda Z, Hiroi NF, Kobayashi TJ, et al. 3D convolutional neural networks-based segmentation to acquire quantitative criteria of the nucleus during mouse embryogenesis. NPJ Syst Biol Appl. 2020;6:32.

14. Hollandi R, Szkalisity A, Toth T, Tasnadi E, Molnar C, Mathe B, et al. nucleAIzer: A Parameter-free Deep Learning Framework for Nucleus Segmentation Using Image Style Transfer. cels. 2020;10:453–8.e6.

15. Stringer C, Wang T, Michaelos M, Pachitariu M. Cellpose: a generalist algorithm for cellular segmentation. Nat Methods. 2020. doi: 10.1038/s41592-020-01018-x.

16. Hollandi R, Diósdi Á, Hollandi G, Moshkov N, Horváth P. AnnotatorJ: an ImageJ plugin to ease hand annotation of cellular compartments. Mol Biol Cell. 2020;31:2179–86.

17. Tasnadi EA, Toth T, Kovacs M, Diosdi A, Pampaloni F, Molnar J, et al. 3D-Cell-Annotator: an open-source active surface tool for single-cell segmentation in 3D microscopy images. Bioinformatics. 2020;36:2948–9.

18. Berg S, Kutra D, Kroeger T, Straehle CN, Kausler BX, Haubold C, et al. ilastik: interactive machine learning for (bio)image analysis. Nat Methods. 2019;16:1226–32.

19. Bazin P-L, Cuzzocreo JL, Yassa MA, Gandler W, McAuliffe MJ, Bassett SS, et al. Volumetric neuroimage analysis extensions for the MIPAV software package. J Neurosci Methods. 2007;165:111–21.

20. Shattuck DW, Leahy RM. BrainSuite: an automated cortical surface identification tool. Med Image Anal. 2002;6:129–42.

21. Wang Y, Li Q, Liu L, Zhou Z, Ruan Z, Kong L, et al. TeraVR empowers precise reconstruction of complete 3-D neuronal morphology in the whole brain. Nat Commun. 2019;10:3474.

22. Bamford P, Lovell B. Unsupervised cell nucleus segmentation with active contours. Signal Processing. 1998;71:203–13.

23. Schroeder W, Martin K, Lorensen B. Visualization Toolkit: An Object-Oriented Approach to 3D Graphics, 4th Edition. 4th edition. Kitware; 2006.

24. The Qt Company. Qt. https://www.qt.io/. xAccessed 4 Dec 2020.

25. Otsu N. A Threshold Selection Method from Gray-Level Histograms. IEEE Trans Syst Man Cybern. 1979;9:62–6.

26. Dice LR. Measures of the amount of ecologic association between species. Ecology. 1945;26:297–302.

27. He K, Gkioxari G, Dollár P, Girshick R. Mask R-CNN. In: 2017 IEEE International Conference on Computer Vision (ICCV). 2017. p. 2980–8.

