## Supplementary material for "Segmentor: A tool for manual refinement of 3D microscopy annotations": Survey Results

### Q1. To what extent do you find the ability to edit voxels in the 2D view useful?

| Field | Minimum | Maximum | Mean | Std Deviation | Count |
|---|---|---|---|---|---|
| Q1. To what extent do you find the ability to edit voxels in the 2D view useful? | 4.00 | 7.00 | 6.33 | 1.11 | 6 |

### Q2. To what extent do you find the ability to edit voxels in the 3D view useful?

| # | Field | Minimum | Maximum | Mean | Std Deviation | Count |
|---|---|---|---|---|---|---|
| 1 | Q2. To what extent do you find the ability to edit voxels in the 3D view useful? | 6.00 | 7.00 | 6.83 | 0.37 | 6 |

### Q3. To what extent do you find the 2D visualizations useful?

| # | Field | Minimum | Maximum | Mean | Std Deviation | Count |
|---|---|---|---|---|---|---|
| 1 | Q3. To what extent do you find the 2D visualizations useful? | 4.00 | 7.00 | 5.50 | 1.26 | 6 |

### Q4. To what extent do you find the 3D visualizations useful?

| # | Field | Minimum | Maximum | Mean | Std Deviation | Count |
|---|---|---|---|---|---|---|
| 1 | Q4. To what extent do you find the 3D visualizations useful? | 7.00 | 7.00 | 7.00 | 0.00 | 6 |

### Q5. To what extent do you find the ability to filter the visibility of segmented regions useful?

| # | Field | Minimum | Maximum | Mean | Std Deviation | Count |
|---|---|---|---|---|---|---|
| 1 | Q5. To what extent do you find the ability to filter the visibility of segmented regions useful? | 5.00 | 7.00 | 6.50 | 0.76 | 6 |

### Q6. To what extent do you find the region splitting feature useful?

| # | Field | Minimum | Maximum | Mean | Std Deviation | Count |
|---|---|---|---|---|---|---|
| 1 | Q6. To what extent do you find the region splitting feature useful? | 6.00 | 7.00 | 6.67 | 0.47 | 6 |

### Q7. To what extent do you find the region growing/shrinking feature useful?

| # | Field | Minimum | Maximum | Mean | Std Deviation | Count |
|---|---|---|---|---|---|---|
| 1 | Q7. To what extent do you find the region growing/shrinking feature useful? | 4.00 | 7.00 | 6.33 | 1.11 | 6 |

### Q8. To what extent do you find the region merging feature useful?

| # | Field | Minimum | Maximum | Mean | Std Deviation | Count |
|---|---|---|---|---|---|---|
| 1 | Q8. To what extent do you find the region merging feature useful? | 4.00 | 7.00 | 6.00 | 1.41 | 6 |

### Q9. To what extent do you find the synchronized rotation and translation of the 2D and 3D views useful?

| # | Field | Minimum | Maximum | Mean | Std Deviation | Count |
|---|---|---|---|---|---|---|
| 1 | Q9. To what extent do you find the synchronized rotation and translation of the 2D and 3D views useful? | 4.00 | 7.00 | 5.33 | 1.25 | 6 |

### Q10. To what extent do you find the ability to show arbitrary (non-axis-aligned) slices in the 2D view useful?

| # | Field | Minimum | Maximum | Mean | Std Deviation | Count |
|---|---|---|---|---|---|---|
| 1 | Q10. To what extent do you find the ability to show arbitrary (non-axis-aligned) slices in the 2D view useful? | 1.00 | 7.00 | 3.33 | 2.43 | 6 |

### Q11. To what extent do you find the region table useful?

| # | Field | Minimum | Maximum | Mean | Std Deviation | Count |
|---|---|---|---|---|---|---|
| 1 | Q11. To what extent do you find the region table useful? | 6.00 | 7.00 | 6.83 | 0.37 | 6 |

### Q12. To what extent do you find linked region highlighting between the region table and the 3D view useful?

| # | Field | Minimum | Maximum | Mean | Std Deviation | Count |
|---|---|---|---|---|---|---|
| 1 | Q12. To what extent do you find linked region highlighting between the region table and the 3D view useful? | 6.00 | 7.00 | 6.50 | 0.50 | 6 |

### Q13. To what extent do you find linked region selection between the 2D view, 3D view, and region table useful?

| | Field | Minimum | Maximum | Mean | Std Deviation | Count |
|---|---|---|---|---|---|---|
| 1 | Q13. To what extent do you find linked region selection between the 2D view, 3D view, and region table useful? | 5.00 | 7.00 | 6.50 | 0.76 | 6 |

### Q14. To what extent do you find the ability to mark regions as "done" useful?

| # | Field | Minimum | Maximum | Mean | Std Deviation | Count |
|---|---|---|---|---|---|---|
| 1 | Q14. To what extent do you find the ability to mark regions as "done" useful? | 6.00 | 7.00 | 6.67 | 0.47 | 6 |

### Q15. To what extent do you find voxel-level editing (painting, erasing, etc.) useful?

| # | Field | Minimum | Maximum | Mean | Std Deviation | Count |
|---|---|---|---|---|---|---|
| 1 | Q15. To what extent do you find voxel-level editing (painting, erasing, etc.) useful? | 6.00 | 7.00 | 6.83 | 0.37 | 6 |

### Q16. To what extent do you find region-level editing (splitting, merging, etc.) useful?

| # | Field | Minimum | Maximum | Mean | Std Deviation | Count |
|---|---|---|---|---|---|---|
| 1 | Q16. To what extent do you find region-level editing (splitting, merging, etc.) useful? | 6.00 | 7.00 | 6.83 | 0.37 | 6 |

### Q17. To what extent do you feel that the 2D features help increase your segmentation accuracy?

| # | Field | Minimum | Maximum | Mean | Std Deviation | Count |
|---|---|---|---|---|---|---|
| 1 | Q17. To what extent do you feel that the 2D features help increase your segmentation accuracy? | 6.00 | 7.00 | 6.67 | 0.47 | 6 |

### Q18. To what extent do you feel that the 3D features help increase your segmentation accuracy?

| # | Field | Minimum | Maximum | Mean | Std Deviation | Count |
|---|---|---|---|---|---|---|
| 1 | Q18. To what extent do you feel that the 3D features help increase your segmentation accuracy? | 6.00 | 7.00 | 6.83 | 0.37 | 6 |

### Q19. To what extent do you feel that the 2D features help increase your segmentation speed?

| # | Field | Minimum | Maximum | Mean | Std Deviation | Count |
|---|---|---|---|---|---|---|
| 1 | Q19. To what extent do you feel that the 2D features help increase your segmentation speed? | 5.00 | 7.00 | 6.00 | 0.82 | 6 |

### Q20. To what extent do you feel that the 3D features help increase your segmentation speed?

| # | Field | Minimum | Maximum | Mean | Std Deviation | Count |
|---|---|---|---|---|---|---|
| 1 | Q20. To what extent do you feel that the 3D features help increase your segmentation speed? | 5.00 | 7.00 | 6.17 | 0.90 | 6 |

### Q21. To what extent do you feel that the 3D visualizations are useful to give you a greater understanding of the shape of segmented regions vs. the 2D visualizations?

| # | Field | Minimum | Maximum | Mean | Std Deviation | Count |
|---|---|---|---|---|---|---|
| 1 | Q21. To what extent do you feel that the 3D visualizations are useful to give you a greater understanding of the shape of segmented regions vs. the 2D visualizations? | 6.00 | 7.00 | 6.83 | 0.37 | 6 |

### Q22. To what extent do you feel that refining an existing automatic segmentation is useful (vs. starting from nothing)?

| # | Field | Minimum | Maximum | Mean | Std Deviation | Count |
|---|---|---|---|---|---|---|
| 1 | Q22. To what extent do you feel that refining an existing automatic segmentation is useful (vs. starting from nothing)? | 4.00 | 7.00 | 6.00 | 1.15 | 6 |

### Q23. To what extent do you find the separate navigation and editing modes useful?

| # | Field | Minimum | Maximum | Mean | Std Deviation | Count |
|---|---|---|---|---|---|---|
| 1 | Q23. To what extent do you find the separate navigation and editing modes useful? | 5.00 | 7.00 | 6.50 | 0.76 | 6 |

### Q24. Please provide feedback/comments with specific reference to any previous questions as needed (e.g., Question #- insert feedback/comments).

| Q24. Please provide feedback/comments with specific reference to any previous questions as needed (e.g., Question #- insert feedback/comments). |
|---|
| N/A |
| Without being able to go back and reference questions, I realize that referencing specific questions is difficult! The only time I find usefulness in synchronized 2D+3D occurs when I orient to the z-plane with filtering on. Otherwise, I find the non-axis-aligned planes to distract and slow down my tracing speed. |
| n/a |
| N/A |

### Q25. What features did you find the most helpful to improve your tracing speed?

| Q25. What features did you find the most helpful to improve your tracing speed? |
|---|
| 3D feature, merging, splitting |
| increasing the brush size |
| Region splitting and merging are particularly helpful to improve my speed. |
| 3D visualization, filter, the axis planes, and the done checkbox |
| The brush radius tool is especially helpful when removing/creating nuclei. |
| Being able to increase brush radius and splitting. |

### Q26. What features did you find the most helpful to improve your tracing accuracy?

| Q26. What features did you find the most helpful to improve your tracing accuracy? |
|---|
| starting off with the 2D and then looking at the 3D made me improve |
| Rotating the 3D view to resolve the nucleus profile is particularly helpful in identifying voxels (e.g., 'voids' and also 'stray, noncontiguous') I may have missed while tracing in 2D. |
| Editing the cells in 2D before 3D editing is very helpful in visualizing the cell and its position relative to other nuclei for any particular slice. If I were to just use the 3D view for tracing, my cells likely would not be as accurate. |
| Changing the resolution settings and filtering to a single nuclei. |
| 3D,2D |
| 3D visualization, 2D visualization, the axis planes, the splitting feature |

### Q27. Are there any features you would like to add?

| Q27. Are there any features you would like to add? |
| --- |
| N/A |
| N/A |
| I would love to see a 'shadow' boundary of an (n-1) slice (e.g., slice 1 shadow boundary while tracing on slice 2) to help 'build' the nucleus profile. I think this will improve tracing speed for some who like to use arrows keys to check traces between consecutive slices. Also, a 'flood fill' tool, in which I can use the mouse as a 'lasso' to trace the shape on 2D would also save time when 'painting' voxels for a given nucleus. |
| n/a |
| The ability to change the color of the nuclei would be very useful. Editing the colors would allow for grouping, or a better way to visualize cells that are close to each other. In other words, two orange nuclei overlayed closely to each other might be difficult to visualize, so being able to change the color of one of those cells would be incredibly useful. |
| Possibly a timer/stopwatch to keep track of tracing time. A "help center" that gives quick access to quick keys and tips and tricks, rather than having to access the GitHub wiki page. |

### Q28. Are there any other changes you would make?

| Q28. Are there any other changes you would make? |
| --- |
| N/A |
| N/A |
| In thinking about the citizen science platform, one of my concerns is that screen brightness greatly affects the tracer's interpretation of a nucleus boundary. We either need to continue instructing tracers to trace on the highest screen brightness settings (which is a bit of an eye strain), or we have to think of a way to minimize variability in boundary interpretation via thresholding methods. |
| n/a |
| The ability to split nuclei quickly is great, but segmentation times often increase when the cells split in random ways. If there was a way to edit the nuclei prior to the split to ensure that the cells split as close to the desired cute as possible, that would decrease segmentation time and accuracy. |
| For first time users, there should be an automatic tutorial built into the software so that when you open it there are guides pointing you to tools and tell you what to do. |
